# A macaque connectome for large-scale network simulations in TheVirtualBrain

**DOI:** 10.1101/480905

**Authors:** Kelly Shen, Gleb Bezgin, Michael Schirner, Petra Ritter, Stefan Everling, Anthony R. McIntosh

## Abstract

Models of large-scale brain networks that are informed by the underlying anatomical connectivity contribute to our understanding of the mapping between the structure of the brain and its dynamical function. Connectome-based modelling is a promising approach to a more comprehensive understanding of brain function across spatial and temporal scales, but it must be constrained by multi-scale empirical data from animal models. Here we describe the construction of a macaque connectome for whole-cortex simulations in TheVirtualBrain, an open-source simulation platform. We take advantage of available axonal tract-tracing datasets and enhance the existing connectome data using diffusion-based tractography in macaques. We illustrate the utility of the connectome as an extension of TheVirtualBrain by simulating resting-state BOLD-fMRI data and fitting it to empirical resting-state data.

## Background & Summary

Linking cellular and circuit-level neural activity with macroscopic signals collected using noninvasive imaging techniques remains a significant challenge to understanding the underlying mechanisms associated with BOLD-fMRI and M/EEG signals. TheVirtualBrain (TVB) is an open-source software platform developed to meet this challenge through simulations of whole-brain network dynamics constrained by multimodal neuroimaging data^1,2^. TVB models link biophysical parameters at the cellular level with systems-level functional neuroimaging signals^3^. This linkage across spatial and temporal scales needs to be constrained by multi-scale data available from animal models. Recently, simulations of whole mouse brain dynamics have been made available in TVB^4^. In addition to rodents, nonhuman primates have become a valuable animal model for studying large-scale network interactions^5,6^. With unique similarities in brain, cognition, and behaviour to humans^7–9^, nonhuman primate models are well poised to bridge multiple scales of investigation. Here, we describe the development of a macaque structural connectome as an initial step towards establishing a multi-scale macaque model in TVB.

Simulations of large-scale network dynamics in which neural mass models are coupled together depend on neuroanatomical connectivity to define the spatial and temporal interactions of the system^10,11^. The connectome is a critical aspect of these models, with its white matter fiber tract capacities (i.e., weights) and tract lengths acting to scale network interactions. However, the construction of any individual connectome is nontrivial and, in the macaque, can be approached using a variety of datatypes. Neuroanatomical data from axonal tract-tracing studies are commonly considered the gold standard and are publicly available from a few sources. One, for example, is the CoCoMac database^12,13^, which can be queried for any number of cortical and subcortical brain regions^14,15^. It offers extensive coverage of the macaque brain but its description of fiber tract capacities is limited to categorical assignments. In another tracer dataset, Markov and colleagues^16^ provide a fully-weighted description of the macaque connectome. The edge-complete connectome, however, describes only 29 of 91 brain regions of a single hemisphere and its utility for whole brain simulations is limited. Alternatively, connectome construction could be performed using tractography on diffusion weighted imaging (DWI) data, as is commonly done for human subjects^17^. Probabilistic tractography has been shown to yield reasonable estimates of fiber tract capacities in macaques^18–20^ and can also provide estimates of tract length. But the resulting connectomes are undirected and therefore unable to capture the known impact of structural asymmetries on functional network topology^21,22^ and dynamics^23^. Moreover, connectomes derived from probabilistic tractography are tainted by a large number of false positive connections^24,25^ that can dramatically change the topology of the reconstructed network^20,26,27^.

We approached the problem of constructing a fully-weighted whole-cortex macaque connectome by synthesizing the information available from both tracer and tractography data (Fig. 1). We took advantage of the specificity of tracer connectivity and enhanced a whole-cortex tracer connectome with weights estimated from tractography. Consistent with previous studies^28–30^, we have recently shown using a diffusion imaging dataset from macaques that the accuracy of tractography algorithms varies as a function of their parameter settings^20^. Drawing on these findings, we first optimized the tractography algorithm to best reproduce the fully-weighted but partial-cortex tracer connectome from Markov and colleagues^16^ before estimating whole-cortex connectome weights. We also estimated connectome tract lengths using tractography, which for tracer datasets is usually limited to Euclidean or geodesic distance estimates between ROI centers.

**Figure 1.**
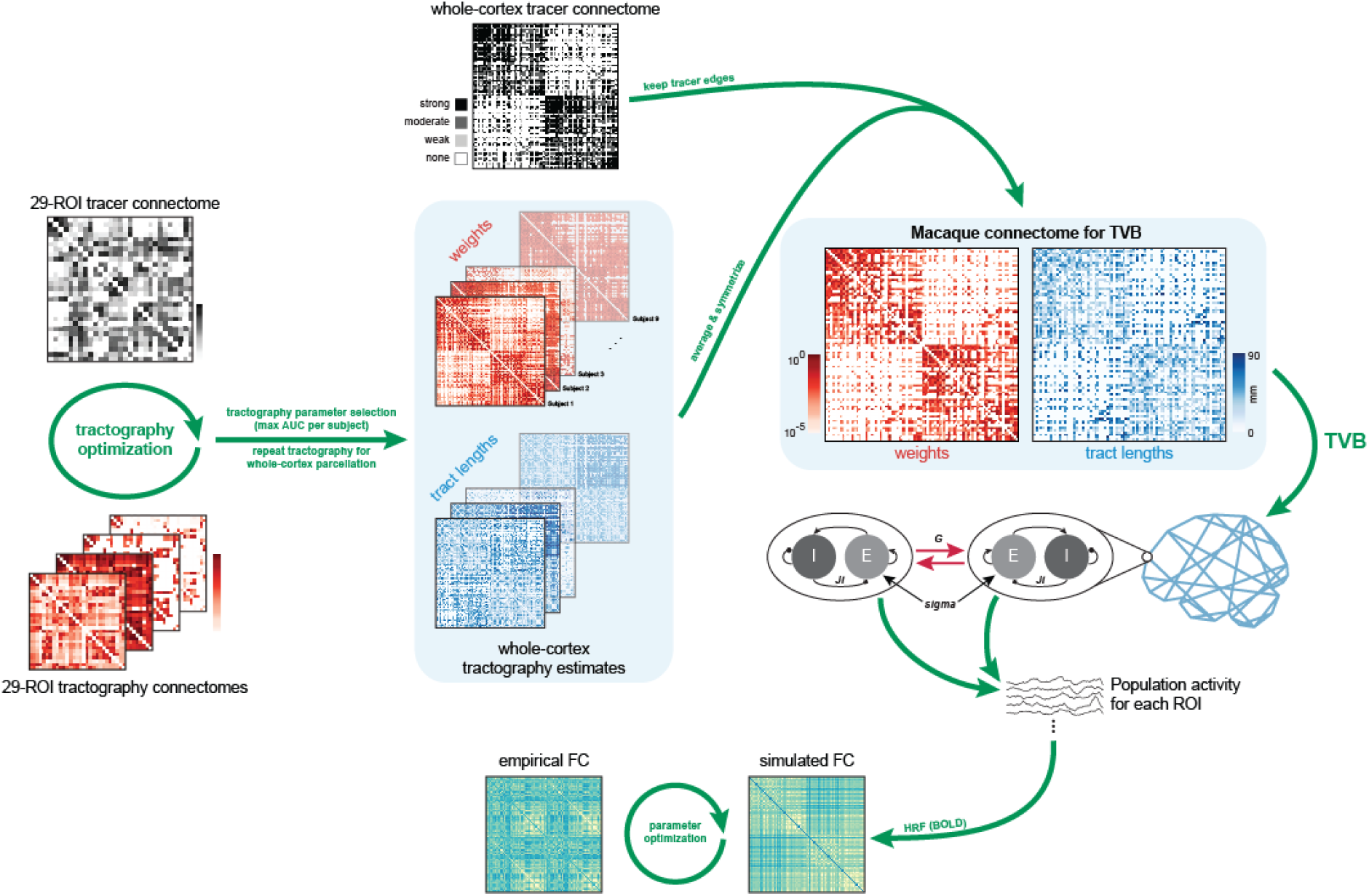
Schematic outline of connectome construction and rsfMRI simulations in TheVirtualBrain (TVB). Tractography parameters were optimized for each subject by comparing tractography results using the 29-ROI parcellation to the 29-ROI tracer connectome from Markov and colleagues^31^ while tractography and connectome construction parameters were varied. The set of parameters yielding the maximum AUC were selected and tractography repeated using the whole-cortex RM parcellation. Tractography-based estimates of weights and tract lengths were averaged and symmetrized and then applied to the nonzero elements of the whole-cortex tracer network. The resultant connectome consisted of a weights and tract lengths matrix, which served as input to TVB for simulations. In this example, the Reduced Wong-Wang-Deco^32^ neural field model was used to simulate population activity for each ROI and a Balloon-Windkessel hemodynamic model^33,34^ was applied to generate BOLD signals. Simulated FC was computed for each parameter set and data fitting to empirical FC was performed to obtain the optimal simulation parameter set.

The connectome presented here will allow us to simulate whole-brain dynamics in the macaque across different modalities and scales (e.g., LFP, M/EEG, fMRI). Simultaneous recordings of neural signals at different spatial and temporal scales in the macaque are already possible^35–37^ and these types of data paired with TVB simulations will allow us to assess our understanding of brain dynamics across scales. In conjunction with existing macaque empirical data and open-access neuroimaging sharing initiatives like PRIME-DE^38^, the virtualized macaque brain will be a solid step forward in our efforts to model structure-function mapping in the whole brain.

## Methods

### Tracer data

Two independent tract-tracing datasets were used in the construction of the whole-cortex macaque connectome. The first was a whole-cortex structural connectivity matrix of 82 regions-of-interest (ROIs) as described in Shen et al^22^. This connectivity matrix follows the Regional Map (RM) parcellation of Kötter and Wanke^39^ and was derived from the CoCoMac database of axonal tract-tracing studies [cocomac.g-node.org]^12–14^. The matrix is directed and includes interhemispheric connectivity. Its edge weights are categorical (‘weak’, ‘moderate’, or ‘strong’).

The second tract-tracing dataset was the edge-complete connectivity matrix of 29 ROIs as described in Markov et al^16^ (Fig. 2a) [data available at core-nets.org]. This connectivity matrix represents intrahemispheric connectivity between a subset of regions in a 91-ROI parcellation. Its edges are fully weighted, spanning six orders of magnitude. This connectivity matrix was used to optimize tractography within the macaque brain. For the purposes of comparison with diffusion-derived connectivity matrices, the tracer matrix was symmetrized by taking the mean of the edge weights of both directions between pairs of ROIs.

**Figure 2.**
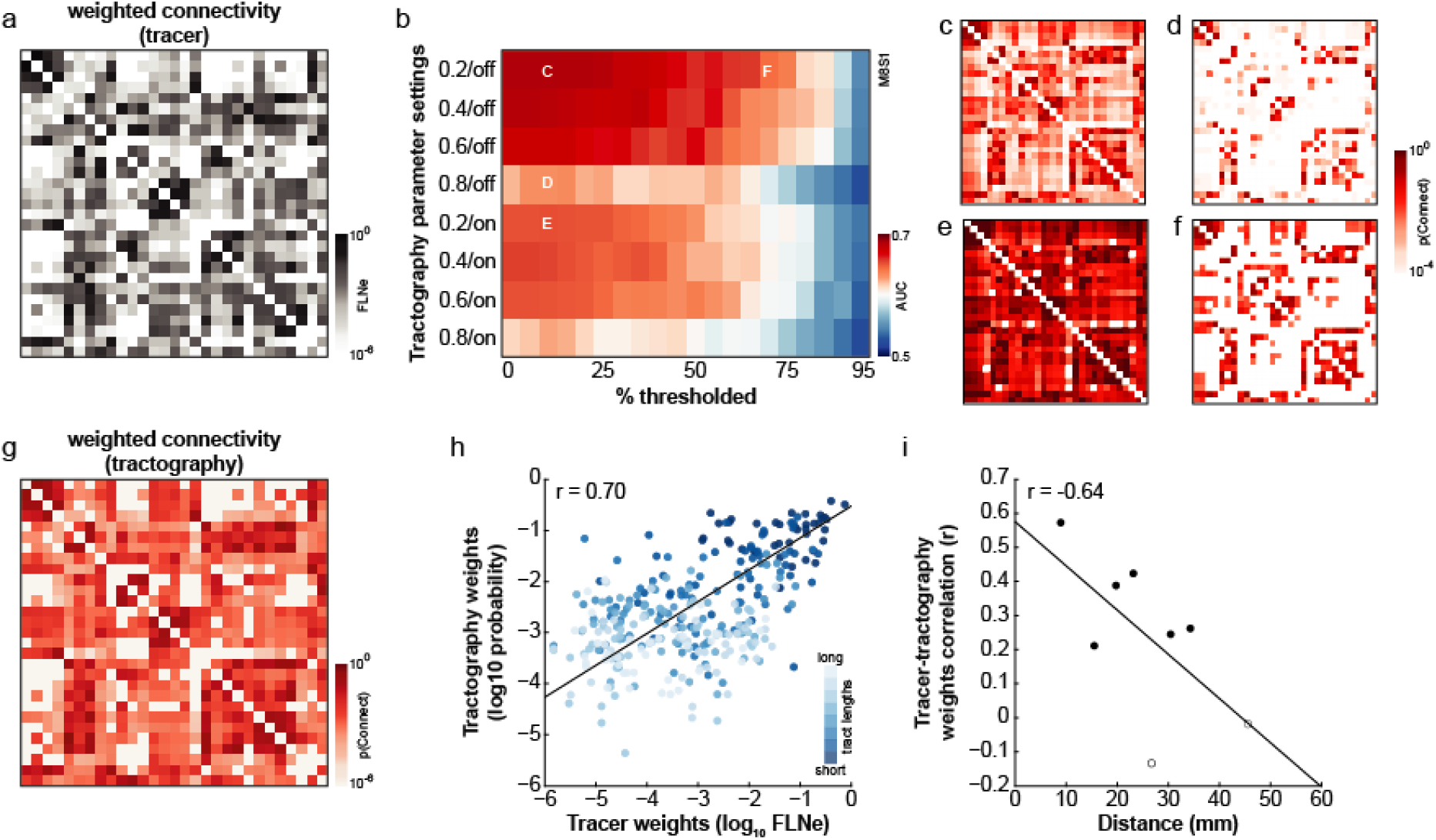
Optimizing connectome construction using existing tracer-derived weights. (a) Tracer-derived 29-ROI weighted connectivity matrix^16^, symmetrized for comparison with tractography-derived matrices. (b) Effect of tractography parameters (y-axis) and connectome thresholding (x-axis) on accuracy of connectome construction for an example subject. Tractography curvature threshold was varied (0.2 – 0.8) and distance correction was toggled ‘on’ or ‘off’. Example matrices from the parameter space as indicated in (c) are shown in panels c-f. Tractography-derived 29-ROI connectivity matrices for this subject showing the (c) most accurate connectome (maximum AUC: 0.69); (d) effect of lowering the curvature threshold (AUC: 0.63); (e) effect of distance correction (AUC: 0.65); and (f) effect of discarding the weakest connections (AUC: 0.64). (g) Tractography-derived weighted connectivity matrix. Only weights for which there is a connection in the tracer connectome are shown. (h) Tractography weight estimates as a function of tracer weights. Connections are colour-coded according to their tract lengths, binned into eight equally-sized bins. Connections belonging to the longest tract length bin denoted in darkest blue with subsequent tract length bins denoted in progressively lighter shades of blue. (i) Pearson correlations coefficients between tracer and tractography weights were negatively correlated with tract length (r = −0.64, bootstrapped CI: −0.98, −0.21). However, even at longer distances, correlation coefficients were statistically significant (p < 0.05, indicated with filled data points).

### ROI parcellations

Volumetric parcellations corresponding to both tracer-based connectivity matrices were defined on the F99 macaque atlas^40^. Both parcellations were registered to each subject’s diffusion data to define ROIs for tractography (see *preprocessing* section below). The RM parcellation was also used for defining ROIs in the resting-state BOLD-fMRI data in order to compute functional connectivity. For the RM parcellation, ROIs were first drawn on the F99 surface^41^ and converted to a labelled volume with a 2 mm extrusion^22^ using the Caret software package^42^ [http://brainvis.wustl.edu/wiki/index.php/Caret:About]. The 91-ROI parcellation and the F99 macaque atlas were both obtained from the SumsDB database [http://brainvis.wustl.edu/wiki/index.php/Sums:About]. Only the 29 ROIs matching the edge-complete tracer matrix were considered and we therefore refer to this parcellation subsequently as the 29-ROI parcellation.

### Neuroimaging data acquisition & preprocessing

Neuroimaging data from 9 male adult macaque monkeys (8 *Macaca mulatta,* 1 *Macaca fascicularis)* were used for the construction and validation of the macaque connectome^43^. These neuroimaging data were a subset of data presented in previous studies^20,44^. The methods described here for diffusion image preprocessing, tractography and connectome construction were developed following the findings of our related work that examined the ability of probabilistic tractography to faithfully reconstruct large-scale network topology^20^. All surgical and experimental procedures were approved by the Animal Use Subcommittee of the University of Western Ontario Council on Animal Care and were in accordance with the Canadian Council of Animal Care guidelines.

Surgical preparation and anaesthesia^45^ as well as imaging acquisition and preprocessing protocols^20,44^ have been previously described. Briefly, animals were lightly anaesthetized before their scanning session and anaesthesia was maintained using 1-1.5% isoflurane during image acquisition. Images were acquired using a 7-T Siemens MAGNETOM head scanner with an in-house designed and manufactured coil optimized for the non-human primate head^46^. Two diffusion weighted scans were acquired for each animal, with each scan having opposite phase encoding in the superior-inferior direction at 1 mm isotropic resolution. For six animals, data were acquired with 2D EPI diffusion while for the remaining three animals, a multiband EPI diffusion sequence was used. In all cases, data were acquired with b = 1000 s/mm^2^, 64 directions, 24 slices. Four resting-state BOLD-fMRI scans were also acquired for each animal using a 2D multiband EPI sequence (600 volumes, TR = 1000 ms, 42 slices, resolution: 1 × 1 × 1.1 mm). Finally, a 3D T1w structural volume was also collected for all animals (128 slices, resolution: 500 μm isotropic).

The Brain Extraction Tool (BET) from the FMRIB Software Library package (FSL v5) was used to extract the brain from skull and soft tissue for all image modalities prior to all other preprocessing steps. Brain extraction of T1w images using BET was suboptimal for all but one animal and brain mask outputs from BET for those animals were manually edited using ITK-SNAP^47^ (www.itksnap.org). Segmentation of the T1w images into three tissue classes was performed using FSL’s FAST tool with bias field correction. Diffusion-weighted images were preprocessed using FSL’s *topup, eddy* and *bedpostx* tools for image distortion correction and modelling of fiber directions. Both parcellations in F99 atlas space were nonlinearly registered to each animal’s T1w structural volume and then linearly registered to diffusion space using the Advanced Normalization Tools (ANTS) software package^48^. All images were visually inspected following brain extraction and registrations to ensure correctness.

FSL’s FEAT toolbox was used for preprocessing the fMRI data, which included motion correction, high-pass filtering, registration, normalization and spatial smoothing (FWHM: 2 mm). Motion in the fMRI data was minimal, with an average framewise displacement across all animals and all scans of 0.015 mm (range: 0.011-0.019 mm). Global white matter and cerebrospinal fluid signals were linearly regressed using AFNI’s (Analysis of Functional NeuroImages) 3dDeconvolve function. The global mean signal was not regressed.

### Tractography optimization for fiber tract capacity estimates

Tractography was performed between all ROIs of the 29-ROI parcellation using FSL’s probtrackx2 function. First, white matter and gray matter voxels that were adjacent to each other were identified in the T1w images. This white-gray matter interface was linearly registered to diffusion space. The white matter voxels of the interface were assigned to ROIs based on their adjacent gray matter ROI assignments. If the adjacent gray matter voxel had no assignment due to inaccuracies in the conversion of the parcellation from surface to volume space, the white matter voxel was given the assignment of the nearest gray matter voxel with an ROI assignment. White matter ROIs were used as seed and target masks. The gray matter voxels adjacent to each seed mask were used as their exclusion mask. As the parcellation was defined in the left hemisphere, an exclusion mask of the right hemisphere was also applied. Tractography parameters were set to 5000 seeds per voxel, a maximum of 2000 steps, and 0.5 mm step length. Paths were terminated if they looped back on themselves and rejected if they passed through an exclusion mask. Tractography was performed repeatedly with various curvature thresholds (0.2, 0.4, 0.6, 0.8) as well as with and without distance correction.

Fiber tract capacity estimates (i.e., ‘weights’) between each ROI pair were taken as the number of streamlines detected between them and dividing by the total number of streamlines that were sent from the seed mask, with the exclusion of those streamlines that were rejected or excluded. A structural connectivity matrix of all ROI pairs was generated with these capacity estimates and symmetrized by taking the mean of the estimates of both directions between ROI pairs. This procedure was repeated for the various tractography settings and resulted in a total of eight connectivity matrices (4 curvature thresholds × 2 distance correction options) for each subject. Each of these eight matrices were then thresholded by discarding the lowest capacity estimates in increments of 5%, resulting in a total of 160 connectivity matrices for each subject.

The optimal set of tractography parameters for each subject was determined by computing the area-under-the ROC curve (AUC) of the 160 matrices in comparison with the “ground truth” tracer-derived connectivity matrix (see Fig. 2b-f for an example subject). The set of tractography parameters yielding the maximum AUC was determined for each subject. In all cases, the maximum AUC value was obtained without distance correction. This was likely because distance correction of all but the longest connections is known to result in lower true positive rates^49^. Applying distance correction to all tract reconstructions regardless of the distance between seed and target ROIs may therefore have been detrimental to most connections. We have also previously shown how applying distance correction results in worse estimates of connection weights than if distance correction is not applied^20^, likely due to the reweighting of connection strengths as a function of the distance between seed and target ROIs (see Fig. 2e for an example). Optimal curvature thresholds ranged between 0.2 and 0.8, while the optimal percentage of weak connections to discard ranged between 0 and 35% (Table 1). Determining tractography parameters this way allowed for tractography to optimally detect the presence and absence of connections for each individual without regard for the accuracy of weight estimates. This method was chosen in favour of explicitly trying to match the tractography weights to the tracer ones (e.g., tractography parameters selected on the basis of having the maximal correlation coefficient between tractography and tracer edge weights) to avoid the variability in tracer weights that cannot been accounted for here. Since different tracer edges were contributed from different animals^16^, and in some cases from a single animal, it is difficult to know to what extent outlier tracer weights exist in the tracer data. If particularly strong or weak weights are poorly represented in the tracer dataset, then they could incorrectly skew the selection of tractography parameters in their favour.

**Table 1.** Optimal tractography parameters for each subject. In all cases, the maximum AUC value was achieved without distance correction.

| Subject | Curvature threshold | Connectome<br>threshold<br>(% discarded) | Max AUC |
| --- | --- | --- | --- |
| 1 | 0.2 | 5 | 0.70 |
| 2 | 0.2 | 15 | 0.68 |
| 3 | 0.8 | 35 | 0.64 |
| 4 | 0.6 | 5 | 0.69 |
| 5 | 0.6 | 25 | 0.66 |
| 6 | 0.2 | 10 | 0.69 |
| 7 | 0.4 | 5 | 0.69 |
| 8 | 0.4 | 0 | 0.73 |
| 9 | 0.6 | 5 | 0.71 |

Tractography was repeated for the RM parcellation using the optimal set of tractography parameters for each subject in the same manner as described above. For intrahemispheric tracking in the RM parcellation, exclusion masks of the opposite hemisphere were applied. Structural connectivity matrices were symmetrized and thresholded to each subject’s optimal threshold for discarding weak weights (Table 1). Due to the relationship between connection weights and distances (see Fig. 2h), thresholding the weakest weights generally corresponded to discarding the longest connections. For the animals whose tractography results were thresholded, the mean discarded connection distance was 50.4 mm (range: 38.9-58.9 mm) and 27 connections were consistently discarded (i.e., thresholded away in at least 6 of 8 animals). The majority of the consistently discarded connections were false positives (70%; 19/27) and the remaining 8 connections were mostly intrahmemispheric ones between visual and frontal/prefrontal cortices (between V2/VACv and M1/FEF/PMC). The weights of these 8 connections may therefore be poorly represented in the resulting macaque connectome as only a few animals contribute to their estimation. However, the impact of this on the overall connectome is likely very minimal given the size of the connectome, which includes 3389 connections.

Tractography between all ROIs of the RM parcellation was then repeated with the distance correction option in probtrackx2 in order to estimate tract lengths. This was done by first dividing the number of streamlines detected between a seed voxel and the target ROI in the distance corrected tractography by the number of streamlines when distance correction was not used. The median value across all seed voxels was taken as the tract length between the seed and target ROIs.

### Macaque connectome construction

Tractography-derived structural connectivity matrices in the RM parcellation were averaged across subjects. These averaged fiber tract capacity estimates were applied to the corresponding nonzero elements of the tracer-derived matrix. The resulting matrix represents a whole-cortex macaque connectome that is both directed (as informed by tracer data) and weighted (as informed by diffusion imaging data) (Fig. 3a)^50^. For technical validation, the weights were fit with a lognormal function and visualized using MATLAB’s Distribution Fitting Tool from its Statistics Toolbox (MATLAB R2011b).

**Figure 3.**
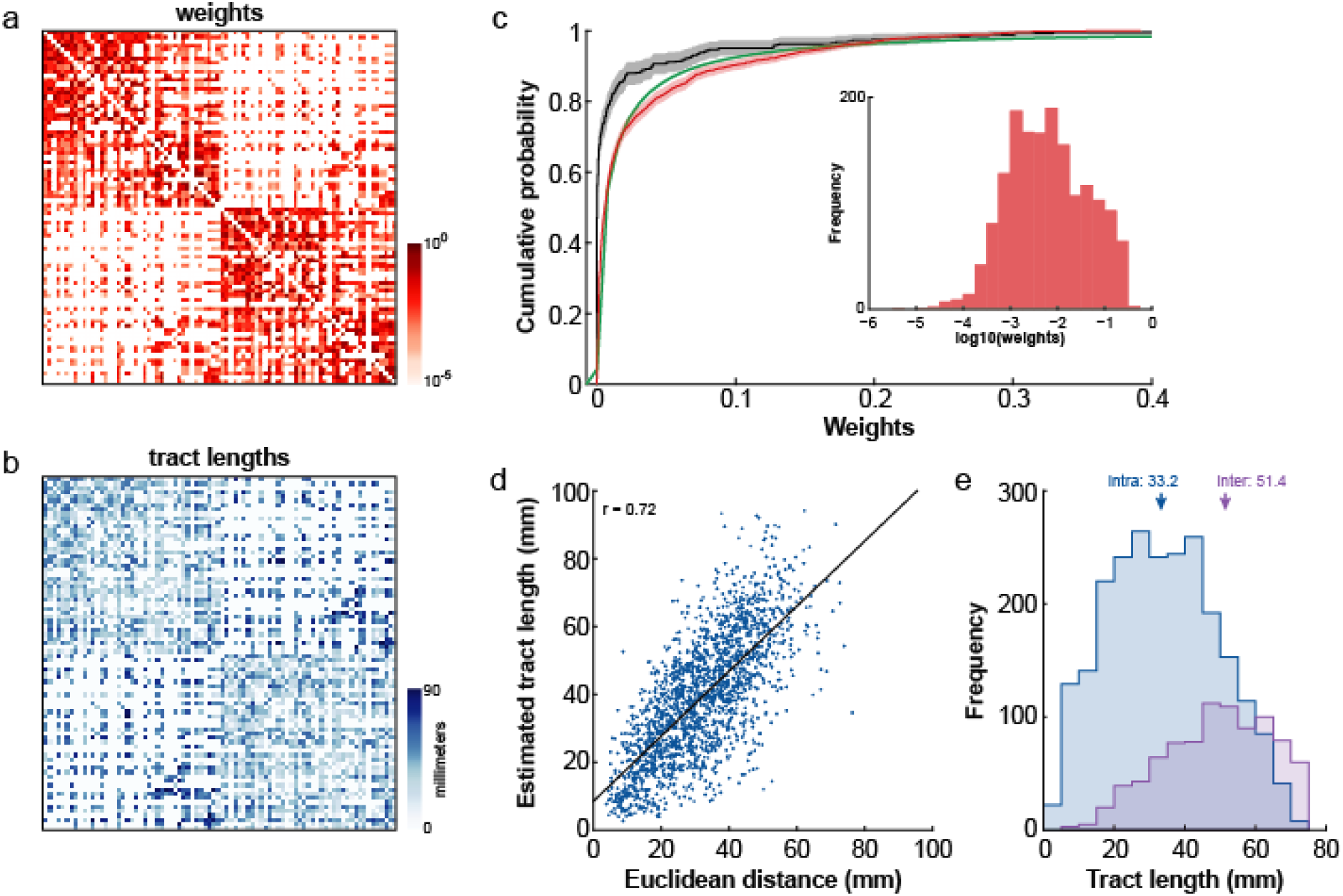
Whole-cortex macaque connectome for large-scale network simulations. Structural connections between 82 ROIs of the RM parcellation were determined from the CoCoMac database of tracer studies. Weights (a) and tract lengths (b) of those connections were estimated using tractography. (c) The lognormal distribution (green) derived from fitting to the tractography-derived weights (red). Tracer weights from the 29-ROI parcellation (cf. Fig. 2a) shown in black. Shading denotes 95% CI. Inset: Frequency distribution of the tractography-derived weights. (d) Relationship between estimated tract lengths and the Euclidean distance between ROI pairs. (e) Frequency distributions of the intra-(blue) and inter- (purple) hemispheric connectivity tract lengths. Median values indicated by arrows.

Tract length estimates were averaged across subjects and the matrix symmetrized. Only those tract lengths with corresponding nonzero elements in the tracer-derived matrix were kept (Fig. 3b)^50^.

### Functional connectivity

Functional connectivity (FC) for each subject was computed by first calculating a weighted average time series^22^ for each ROI in the RM parcellation for each resting-state fMRI scan. The pairwise Pearson correlation coefficients between time series were determined and a Fisher z-transform was applied to each resultant FC matrix. FC matrices were then averaged within subjects to obtain individual FC for each subject. FC matrices were also then averaged across subjects to obtain an overall average FC for the group.

### TVB simulations

Large-scale network dynamics were simulated using TheVirtualBrain (TVB) software platform (thevirtualbrain.org) based on previously described methodologies^32,51,52^. For input to TVB, the connectome’s weights matrix was normalized to its maximum value but not thresholded.

The Reduced Wong-Wang^32^ dynamic mean field model, implemented using C code for efficient parameter exploration, was used to model the local dynamics of each brain region. This mean field model consists of excitatory and inhibitory populations coupled together by excitatory (NMDA) and inhibitory (GABA) synapses. The strength of the coupling from the inhibitory population on the excitatory population was scaled by a feedback inhibition control parameter (*J_i_*). The 82 mean field models, one for each brain region, were then coupled together according to the macaque connectome’s weights matrix and scaled using a global coupling parameter (G). Conduction velocity was fixed across the network so that time delays between regions were governed by the connectome’s tract length matrix. The uncorrelated Gaussian noise of each region was scaled in amplitude by a single noise parameter *(sigma).*

Model fitting was performed by a parameter space exploration in which *G* and *sigma* were varied systematically to optimize the fit between empirical and simulated functional connectivity. These parameters were chosen because previous studies have shown how they are crucial to determining the dynamics of the model^32^ and have been implicated in brain disease^51,52^. *G* was varied between 0.1 and 5 in 100 equally-sized steps while *sigma* was varied between 0.01 and 0.1 in 30 equally-sized steps for a total of 3000 iterations of the parameter search. Excitatory population firing rates have been reported to be in the range of 5 ± 5 Hz in the macaque^53–56^. Tuning the excitatory neural mass models with *J_i_* to constrain their firing rates to ~3 Hz is known to result in better models of FC with biologically-plausible dynamical properties^32^. Therefore, within each parameter search iteration, *J_i_* was first tuned for each region individually to constrain its excitatory population firing rate to ~3 Hz^32^. *Ji* tuning was performed for up to 100 iterations as described by Schirner and colleagues^57^. Simulations in which 100 iterations of *J_i_* tuning still resulted in unrealistic excitatory population firing rates (mean >= 10 Hz across regions) were not considered for analysis. Conduction velocities in the macaque brain have been estimated to be in the range of 3-10 m/s^58,59^ and were set to 4 m/s in our simulations to reflect the median of the distribution reported by Girard and colleagues^58^. All other parameters of the model were the same as those from Deco and colleagues^32^. Resting-state BOLD-fMRI data with the same duration (10 mins) and sampling rate (TR = 1 s) as the empirical BOLD-fMRI data were then simulated by applying a Balloon-Windkessel hemodynamic model^33,34^ to the excitatory synaptic activity of each region. Simulated FC was computed and the cosine similarity (aka uncentered Pearson correlation^32^) between the upper triangles of the simulated and empirical FC matrices was used to determine the goodness of fit. The simulated data with values of *G* and *sigma* giving the maximum cosine similarity to empirical FC was used for further analysis. The statistical significance of our goodness of fit measure was assessed by creating a distribution of 1000 shuffled networks using the empirical FC. This was done by randomly rewiring the empirical network using the ‘null_model_und_sign’ function from the Brain Connectivity Toolbox (BCT; https://sites.google.com/site/bctnet/), which preserved the degree-, weight- and strength-distributions from the empirical network. The cosine similarity between each of the 1000 shuffled networks and the simulated FC was then computed to create a null distribution of cosine similarity values. A p-value was computed as the proportion of the null distribution with a cosine similarity value greater than or equal to the observed value.

A principal component analysis (PCA) was performed on the simulated timeseries data to identify the dominant networks in the simulated resting-state data. The first three components were chosen for interrogation as together they explained 79.1% of the covariance in the data. We arbitrarily selected the 10 nodes with the highest loadings within each component for the purpose of visualization.

As the best fit simulated network had only positively-weighted edges, comparison of its topology with that of the empirical network was done by first thresholding the empirical network to only the positive weights using the BCT function ‘threshold_absolute’ and then thresholding the simulated network to have the same density as the empirical one using the BCT function ‘threshold_proportional’. The sum of each node’s weights within each network were then calculated using the ‘strengths_und’ BCT function to derive the nodal strengths for comparison.

To test the extent to which our average macaque connectome can be used for modelling any individual macaque’s brain dynamics, we repeated our fitting procedure using the empirical FC matrix from each individual subject. To assess the goodness of fit for each subject, we repeated the shuffling procedure on each individual FC matrix to produce nine sets of 1000 shuffled networks. We additionally tested the extent to which using an individual’s structural connectome for simulations could provide a better fit to that individual’s empirical FC by repeating the entire simulation procedure using each animal’s optimized structural connectome (based solely on DWI tractography). As before, weights matrices were normalized to their maximum value but not thresholded.

Significant testing for correlations was done using permutation tests, whereby the the correlation coefficient between two variables *x* and *y* was recalculated after randomly shuffling *y* 1000 times to produce a null distribution of correlation coefficients. The p-value was taken as the proportion of the null distribution with a correlation coefficient greater than or equal to the observed coefficient.

### Code availability

Custom code written in MATLAB for the generation of the macaque connectome, including wrapper scripts for neuroimaging preprocessing and DWI tractography, is available upon request from the corresponding author. The C code for the Reduced Wong-Wang model in TVB is available at https://github.com/BrainModes/TVB_C.

## Data Records

The neuroimaging data used for generating the macaque connectome (T1w & DWI) and for data fitting (fMRI) of the TVB simulations is available at OpenNEURO^43^. Each subject directory contains raw data (reconstructed into NIFTI format from DICOMS but otherwise unprocessed) that includes:

- /anat/

∘ *_T1w.nii.gz: 3D structural volume used for segmentation (GM, WM & CSF) and registration
- /dwi/

∘ *_run-01_dwi.nii.gz, *_run-02_dwi.nii.gz: diffusion weighted images having opposite phase encoding;
∘ *_run-01_dwi.bvec, *_run-01_dwi.bval, *_run-02_dwi.bvec, *_run-01_dwi.bval: corresponding bvec and bval files for DWI images
- /func/

∘ *_task-rest_run-01_bold.nii.gz, *_task-rest_run-02_bold.nii.gz, *_task-rest_run-03_bold.nii.gz, *_task-rest_run-04_bold.nii.gz: four resting-state fMRI scans

In addition, preprocessing derivatives for each subject are provided and include:

- /anat/

∘ *_space-subject_desk-skullstripped_T1w.nii.gz: skull-stripped T1w image
- /dwi/

∘ *_space-subject_desc-eddy_dwi.nii.gz: eddy-corrected DWI images, runs 01 and 02 concatenated
∘ *_space-subject_desc-eddy_dwi.bvec, *_space-subject_desc-eddy_dwi.bval,: corresponding bvec and bval files for eddy-corrected concatenated DWI images
- /func/

∘ *_task-rest_run-01_space-F99_desc-confoundregressed_bold.nii.gz, *_task-rest_run-02_space-F99_desc-confoundregressed_bold.nii.gz, *_task-rest_run-03_space-F99_desc-confoundregressed_bold.nii.gz, *_task-rest_run-04_space-F99_desc-confoundregressed_bold.nii.gz: four preprocessed (up to confound regression) resting-state fMRI scans, registered to F99 macaque template

The macaque connectome dataset is openly available for download from Zenodo^50^. Available files for download include:

- weights.txt: Weights matrix representing a directed and weighted connectome in the 82-node RM whole-cortex parcellation; ROI order as presented in the look-up table
- tract_lengths.txt: Tract length matrix, in mm
- TVBmacaque_RM_LUT.txt: Look-up table for RM parcellation; includes ROI ordering, label indices, ROI names and abbreviations
- RM_inF99.nii.gz: labelled volume of gray matter ROIs of RM parcellation, in F99 macaque atlas space; labels as specified in the look-up table
- surf_faces.txt, surf_vertices.txt, surf_labels.txt: F99 macaque atlas surface and corresponding RM parcellation labels; labels as specified in the look-up table

### Technical Validation

Tractography was used to estimate the fiber tract capacities of the macaque connectome via an optimization procedure that selected the tractography and connectome threshold parameters that produced the most accurate connectome for each subject, as assessed by comparison with tracer data (Figure 2). The average 29-ROI connectome that resulted from this optimization procedure had weights that were correlated with the tracer weights (r = 0.70, p < 0.001)^also see 20^, suggesting that the optimized tractography parameters for each subject could reasonably be applied to the estimation of fiber tract capacities across the whole brain.

Tractography was therefore repeated using the whole-cortex RM parcellation and the average weights and tract length estimates across subjects were applied to the tracer-derived RM connectivity matrix to generate a canonical macaque connectome (Fig 2a-b). As reported in tracer studies, macaque fiber tract capacities follow a lognormal distribution^16,31^. We fit our tractography-derived weight estimates with a lognormal function (mu=-5.07, sigma=1.92) and found that they also generally followed a lognormal distribution (Fig 3c). In sum, despite optimizing tractography parameters to the detection of the presence and absence of connections while ignoring the effect of parameter choices on weight estimates, tractography produced reasonably good estimates of edge weights.

As expected, tract length estimates were correlated with the Euclidean distance between ROI pairs (Pearson correlation, r = 0.72, p < 0.001; Fig. 3d), with tract length estimates (median: 38.8 mm) being on average longer than Euclidean distances (median: 31.4 mm; Wilcoxon rank sum test, p < 0. 001). Tract length estimates were also significantly shorter on average for intrahemispheric connections (median: 33.2 mm) than interhemispheric ones (median: 51.4 mm; Wilcoxon rank sum test, p < 0.001) (Fig 3e). Using a tracer connectome as the basis for the macaque brain’s neuroanatomical pathways is still limited by the lack of available interhemispheric connectivity data as well as by sources of variability in tracer data that have not been considered here (see ^20^ for a discussion). However, it has been shown that limiting false positives at the expense of false negatives allows for better representation of network topology^27^. As such, we believe that building the macaque connectome based on the available tracer data and only using tractography to estimate the edge weights is the best approach given the current limitations of probabilistic tractography.

For the purposes of validating the connectome and demonstrating a use case for the dataset, we simulated macaque resting-state BOLD-fMRI data in TVB. A parameter space exploration that fit the simulated FC to the overall grand average empirical FC across a range of *G* and *sigma* values was performed. The best fit simulated FC occurred with parameter values *G* = 0.298 and *sigma* = 0.0255 (Fig. 4a-c). This corresponded with a cosine similarity fit of 0.75, which was significantly greater than the cosine similarity fits with null networks (p < 0.001). This best fit simulation was chosen for further analysis. Example simulated BOLD timeseries from five randomly chosen ROIs are shown in Figure 4d. We inspected the top 10 nodes from the three most dominant networks identified using a PCA on the simulated BOLD timeseries. The first network (59.3% of the covariance) was comprised of lateral prefrontal and anterior cingulate regions bilaterally as well as left intraparietal area (Fig 4d, red). The second network (12.3% of the covariance) included superior and central temporal cortical regions, primary and secondary auditory cortices, as well as the insula bilaterally (Fig 4e, green). The third (7.6% of the covariance) was composed of V1, V2 and exstrastriate visual cortices bilaterally (Fig 4e, blue). These networks are similar to the frontal, superior temporal, and visual resting-state networks previously reported in macaques^45,60,61^. We additionally computed the nodal degree (sum of weights, or strengths) of the simulated and empirical FC networks to compare their topologies. The strength of each node in the simulated network was correlated with its strength in the empirical network (r = 0.51, p < 0.001) (Fig. 4f).

**Figure 4.**
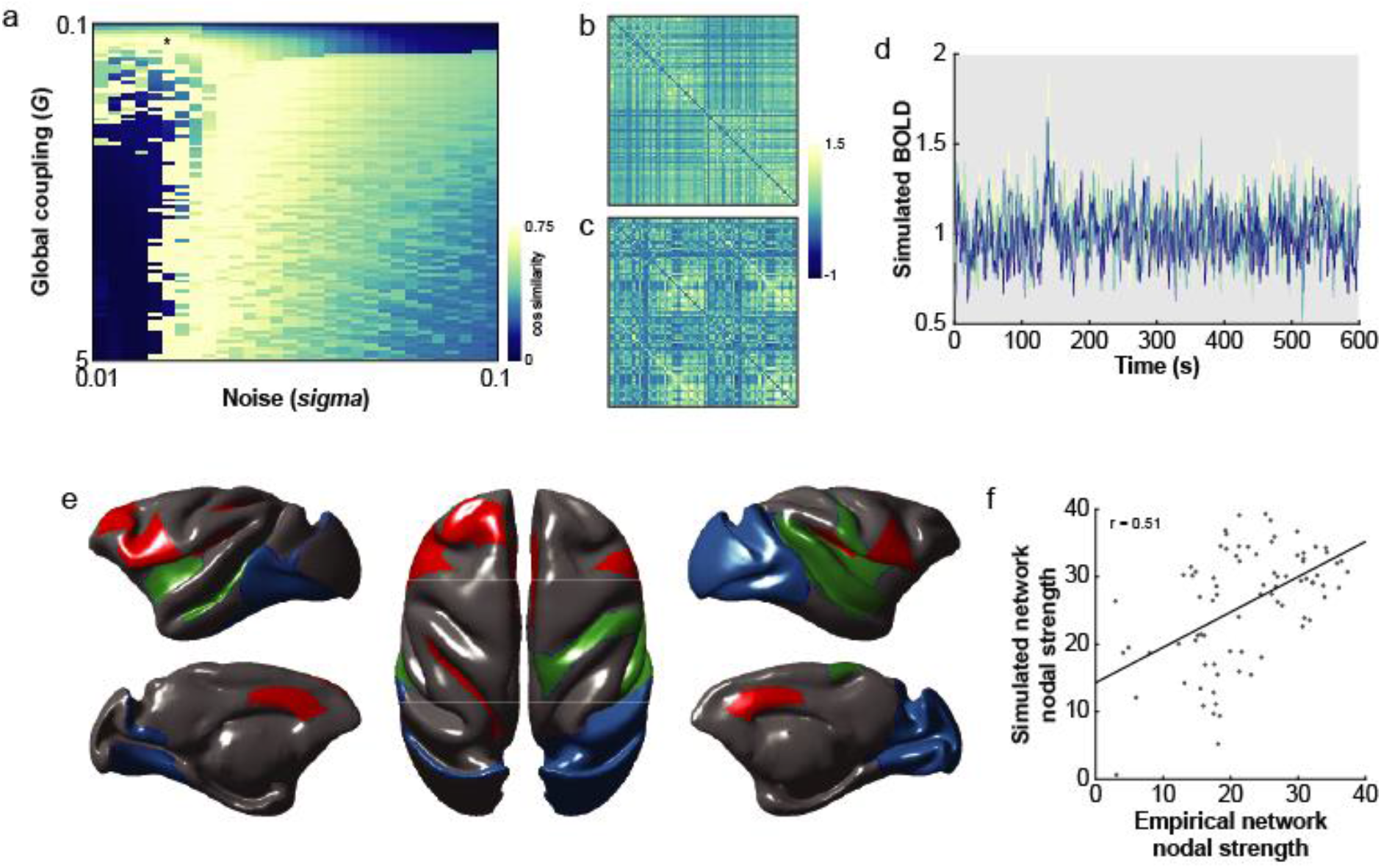
Large-scale network simulations of resting-state fMRI data in the macaque. (a) Goodness of fit (cosine similarity) between simulated FC grand average empirical FC with various instantiations of global coupling *‘G’* and noise *‘sigma’* parameters. Maximum cosine similarity denoted by black asterisk. (b) Simulated FC matrix associated with best fit *G* and *sigma* values. (c) Grand average empirical FC matrix. (d) Simulated BOLD signals of five randomly selected ROIs. (e) Networks associated with the first three principal components (PC) of the simulated timeseries. The 10 nodes having the largest loadings were plotted for the first PC in red, the second PC in green and the third PC in blue. (f) Strength of each node (i.e., sum of its weights) in the simulated network as a function of its strength in the empirical network. Only positive empirical FC was considered, and the simulated network was thresholded to the same density as the average empirical one.

To determine whether a canonical macaque connectome could be used to simulate resting-state BOLD-fMRI of individual macaques, we fit the set of 3000 simulations generated from the parameter space search to the empirical FC of each of our subjects. The best fit cosine similarity for each animal (range: 0.47 – 0.80) (Fig. 5a) was significantly greater than the fits with each animal’s null FC distribution (all p < 0.01). Using each subject’s individual connectome for simulations resulted in similar fits to individual empirical FC (cosine similarity: 0.47 – 0.79) (Fig. 5a) that were not significantly different from the fits obtained using the canonical connectome (paired t-test, p = 0.35). Together with the observation that structural connectomes exhibit far less variance than functional ones^62^, this suggests that a canonical macaque connectome can be used in simulations to capture some aspects of individual macaque whole-brain dynamics.

**Figure 5.**
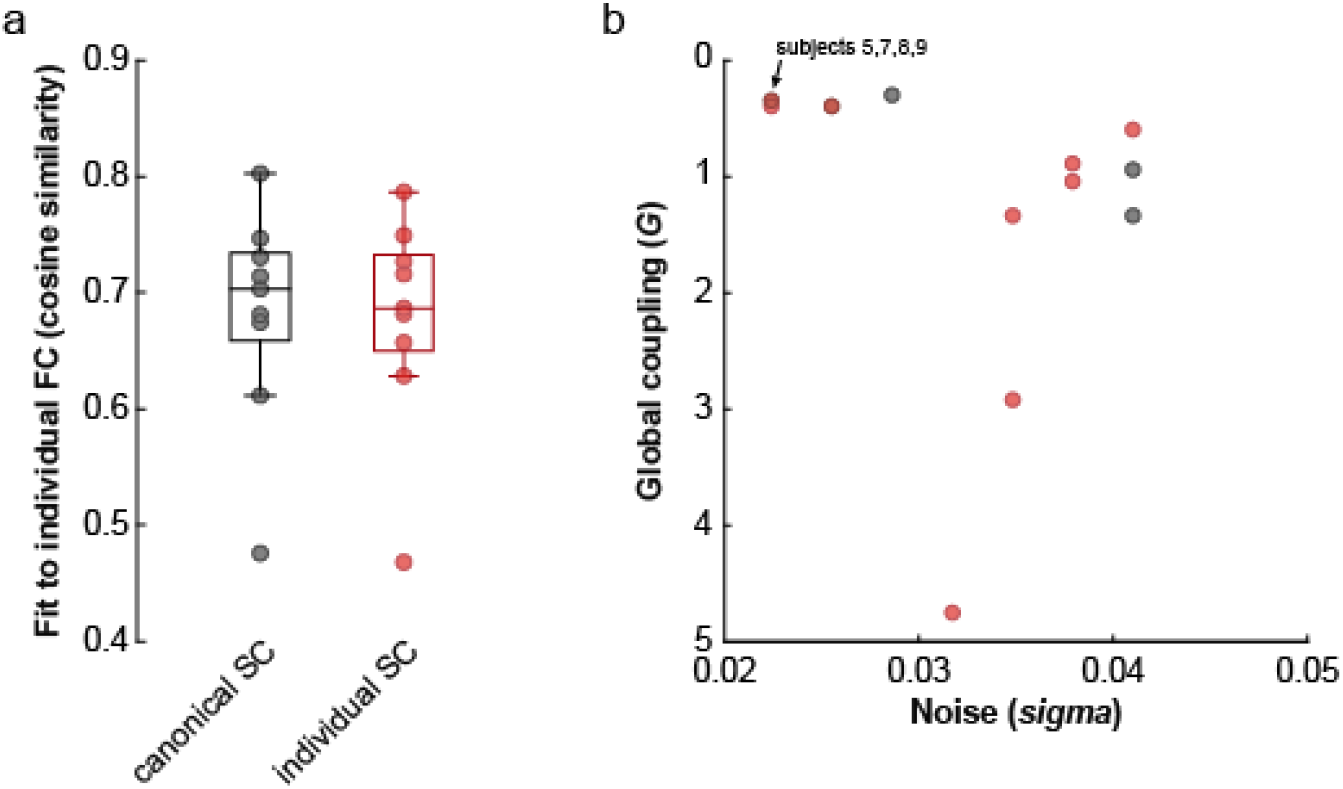
Specificity of TVB simulations when using canonical and individualized connectomes. (a) Goodness of fit (cosine similarity) between simulated FC with individual FC for canonical (black) and individualized (red) connectome simulations. (b) Individual parameter sets shown in parameter space. Using the canonical connectome for simulating individual FC resulted in identical parameter sets for 4 of 9 subjects (black) while individualized connectome simulations gave unique parameter sets (red) for all subjects.

While the fits of the individualized simulations to empirical data may be no better than when using the canonical connectome, an examination of the best-fit simulation parameters under each condition suggests that individual specificity may instead be captured within the biophysical parameters of the model (Fig. 5b). In the case of the canonical connectome simulations, 4 of 9 subjects had identical *G* and *sigma* parameters and the simulated FC were all fairly similar to each other (cosine similarity range: 0.80 – 1). For individualized connectome simulations, however, all parameter sets were different across subjects and resulted in simulated FC that varied more across subjects (cosine similarity range: 0.76 – 0.99). This is in line with our recent findings that individualized TVB model parameters are better at predicting cognitive status in dementia than the coupling between structural and functional connectivity^51^ and suggests that generative models based on individual connectomes may be able to capture subject-specific characteristics that are not identifiable from the empirical data alone. The modelling approach may therefore extract the portion of the measured variance in the empirical data that is most closely related to the underlying biophysics, increasing the sensitivity to individual differences.

### Usage Notes

In the technical validation section above, we described an example of how the macaque connectome dataset can be used for region-based simulations. However, the dataset also includes a labelled cortical surface, allowing for surface-based simulations such as those presented by Proix and colleagues^63^. Further information on the TVB software platform, including the download package, documentation and tutorials can be found at https://www.thevirtualbrain.org (also see https://github.com/the-virtual-brain).

Our choice to construct the macaque connectome using the RM parcellation was a deliberate one. The RM is one of the few whole cortex macaque parcellations whose ROIs are generally well represented by tracer data in CoCoMac^64^. Finer parcellations of the macaque brain such as the full 91-ROI Markov-Kennedy^16^ or the 175-ROI Paxinos^65^ parcellations could be considered in the future in combination with methods such as tractography or distance-based models for estimating missing tracer data^20,66,67^. An additional advantage to the RM parcellation is that it was intended to harmonize cytoarchitectonic, topographic and functional definitions of brain regions across primate species^39^. The application of the RM parcellation to the human brain therefore allows for direct cross-species comparisons. The RM parcellation, with the addition of 14 subcortical regions, and the corresponding connectome in humans is already available in TVB^66^.

## Acknowledgements

This work was supported by grants from the Canadian Institutes of Health Research (CIHR; FRN 148365) and the Canada First Research Excellence Fund (to BrainsCAN) to S.E. P.R was funded by: H2020 Research and Innovation Action grants 826421 and 785907 and ERC 683049; German Research Foundation CRC 1315 and grant RI 2073/6-1; Berlin Institute of Health & Foundation Charité, Johanna Quandt Excellence Initiative. We thank Amanda Easson, Tyler Good, John Griffiths, Zheng Wang and Marmaduke Woodman for technical advice.

## Author contributions

K.S. and A.R.M. designed the study. G.B. and S.E. collected the data. M.S. and P.R. developed the c-code for simulations. K.S. analysed the data and prepared the first draft of this manuscript. All authors contributed to the subsequent revisions of the manuscript.

